# On expert curation and sustainability: UniProtKB/Swiss-Prot as a case study

**DOI:** 10.1101/094011

**Authors:** Sylvain Poux, Cecilia N. Arighi, Michele Magrane, Alex Bateman, Chih-Hsuan Wei, Zhiyong Lu, Emmanuel Boutet, Hema Bye-A-Jee, Maria Livia Famiglietti, Bernd Roechert The UniProt Consortium, The UniProt Consortium

## Abstract

**MOTIVATION:** Biological knowledgebases, such as UniProtKB/Swiss-Prot, constitute an essential component of daily scientific research by offering distilled, summarized, and computable knowledge extracted from the literature by expert curators. While knowledgebases play an increasingly important role in the scientific community, the question of their sustainability is raised due to the growth of biomedical literature.

**RESULTS:** By using UniProtKB/Swiss-Prot as a case study, we address this question by using different literature triage approaches. With the assistance of the PubTator text-mining tool, we tagged more than 10,000 articles to assess the ratio of papers relevant for curation. We first show that curators read and evaluate many more papers than they curate, and that measuring the number of curated publications is insufficient to provide a complete picture. We show that a large fraction of published papers found in PubMed is not relevant for curation in UniProtKB/Swiss-Prot and demonstrate that, despite appearances, expert curation is sustainable.

**AVAILABILITY:** UniProt is freely available at http://www.uniprot.org/.

## Introduction

Biological knowledgebases have become indispensable for biomedical research by providing data in easily accessible formats. The Universal Protein Resource (UniProt) is one such key resource that acts as a central hub of protein knowledge by offering a unified view of protein sequence and functional information (UniProt, 2015). Expert curation constitutes a core activity of the UniProt Knowledgebase (UniProtKB) which is composed of two sections, UniProtKB/Swiss-Prot, the reviewed section containing expertly curated records with information extracted from the literature and curator-evaluated computational analysis, and UniProtKB/TrEMBL, the unreviewed section with computationally analyzed records, enriched with automatic annotation.

Bioinformatic predictions of protein function rely upon correctly annotated database sequences, and the presence of inaccurately or poorly annotated records introduces noise and bias to biological analyses (Bengtsson-Palme, Boulund, et al., 2016). Literature-based expertly curated data is highly reliable, and therefore considered the gold-standard, providing, in the case of UniProtKB, high-quality annotations for experimentally characterized proteins across diverse protein families. In this way, it serves as a source of annotations that can be used for the development and enhancement of bioinformatics algorithms and text mining methods. In addition, literature-based annotations of characterized proteins are the basis for the automatic annotation of uncharacterized ones, a key challenge in the big data era which is witnessing the generation of large amounts of sequences (Oliver, Lock, et al., 2016; Pedruzzi, Rivoire, et al., 2015; UniProt, 2015).

Despite the aforementioned needs and usage of expert curation, the question about its long-term sustainability has frequently been raised. Expert curation is considered to be a time-demanding and expensive activity (Bourne, Lorsch, et al., 2015), especially in light of the continuing growth of the biomedical literature with over 1 million papers published every year. In addition, the number of articles fully curated per year in UniProtKB/Swiss-Prot (between 8,000 and 10,000 articles per year based on the last 7 years) seems low in comparison to the amount of literature available, giving the impression that literature curation cannot scale in the face of the increasing amount of published papers. This picture is however misleading because only publications that provide relevant information are included in UniProtKB/Swiss-Prot and many articles that have been examined during the curation process are not included. However, we have not tracked this information in a formal manner until now.

To give a clearer picture of the landscape of curatable articles and to address concerns about the sustainability of literature-based curation, we performed a study of the ratio of relevant versus non-relevant papers. For this purpose, we used the PubTator text mining system (Wei, Kao, et al., 2013a) to classify articles evaluated during the curation process. We monitored the literature triage process during a 6-month period with a set of curators in order to (i) determine the total number of articles that we read and/or evaluate, (ii) quantify the fraction of the relevant literature covered by UniProtKB and (iii) address the question of the sustainability of expert curation in UniProtKB/Swiss-Prot.

## Methods

### PubTator

PubTator (http://www.ncbi.nlm.nih.gov/bionlp/pubtator) is a web-based application that automatically annotates all 26-million articles in PubMed with key biological concepts via advanced text mining software tools. These include GNormPlus (Wei, Kao, et al., 2015) for identifying genes, tmChem (Leaman, Wei, et al., 2015) for chemicals, DNorm for (Leaman, Dğgan, et al., 2013) diseases, SR4GN (Wei, Kao, et al., 2012) for organisms, and tmVar (Wei, Harris, et al., 2013b) for mutations. PubTator has been widely used for text mining, data curation, and bioinformatics research (e.g. (Singhal, Simmons, et al., 2016; Wei, Leaman, et al., 2016)).

#### Pubtator customizations for UniProt curation

To meet the specific needs of UniProt curation, a number of customizations were made to both the annotation results and user interface. First, all text-mined gene/protein annotations with corresponding NCBI Gene identifiers were converted to UniProt accessions (e.g., NCBI Gene ID: 818982 to UniProtKB accession Q39026). Next, we developed a frequency-based approach for ranking articles with rich protein information. The user interface updates mainly concerned the PubTator’s curation page. We first added a third category for UniProt curators to classify an article – Not priority – in addition to ‘Curatable’ and ‘Not curatable’. Furthermore, five sub-categories were inserted under the existing ‘Not curatable’ category: “Out of scope”, “Redundant”, “High-throughput”, “Insufficient evidence”, and “Review/comment” (Figure 1). Finally, we added a text box to allow users to record their comments.

**Figure 1.**
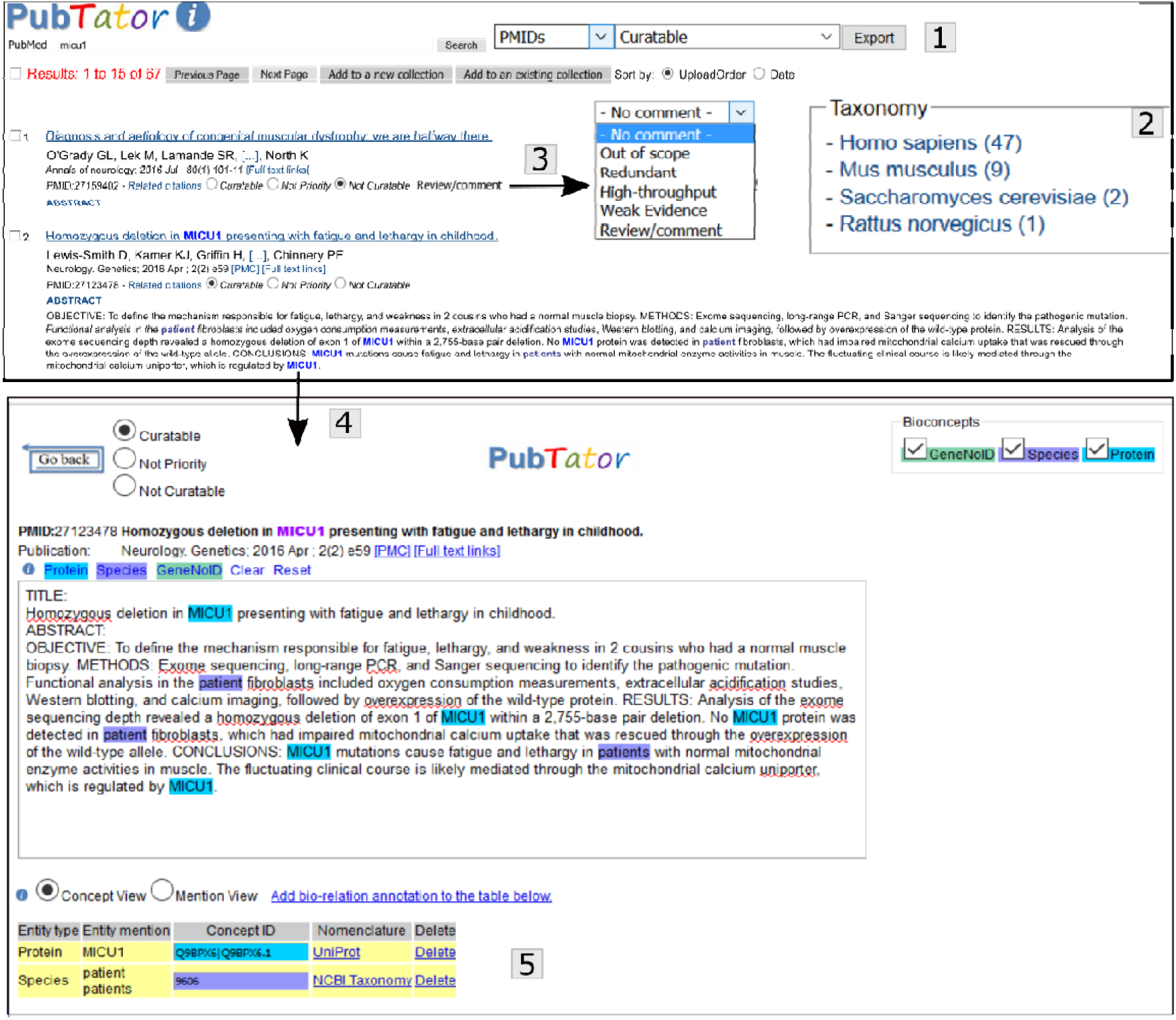
Screenshot of the PubTator tool. Some of PubTator’s functionalities include: (1) export of PubMed identifiers (PMIDs) and annotations for the different sets (e.g., curatable and not-curatable); (2) filtering of articles by species; (3) menu for not-curatable options; (4) access to abstract with annotations; and (5) table with annotations and with links to UniProt accessions.

### Preparation of Datasets

#### Random sampling of 500 PubMed articles

To evaluate the proportion of PubMed articles that are relevant for UniProt curation, we first generated a set of 500 PubMed articles published from 2013 to 2015 (166 articles in 2013, and 167 in both 2014 and 2015) by random sampling. We then applied PubTator to automatically mine protein and organism mentions in this evaluation set.

#### Distribution of article types in PubMed versus random sampling

To check if the distribution of article types (such as news, biography, and journal article) in random sampling was representative of that of PubMed, we accessed PubMed on Sept. 07, 2016, retrieved and recorded the different article types for all (using as query ‘all[filter]’) and for the 500 PubMed random set (by entering the list of PubMed identifiers). In both cases we used the filters for article type (e.g., “biography”[Publication Type]) to record the number in the given category.

#### Weekly collection from selected journals

Each week, PubTator generates an update for new articles published in a selected set of relevant journals for protein research (Cell, Developmental Cell, Elife, Genes and Development, Molecular Cell, Nature Cell Biology, Nature Genetics, Nature, PLoS Biology, PLoS Genetics, Science, The EMBO Journal, The Plant Cell). All new articles are first mined for protein and species information and then ranked based on protein frequency. To make it easy for the curators, the entire set of new articles can be viewed as a whole or separately by each individual journal.

#### Protein-centric curation workflow

The five UniProt curators selected for this analysis work in different annotation programs (for a list of the different annotation programs in UniProtKB/Swiss-Prot, see http://www.uniprot.org/help/?fil=section:biocuration). E.B. is specialized in curation of plant proteins; H.B.-A.-J. is specialized in curation of *Caenorhabditis elegans* proteins; M.L.F. is specialized in curation of proteins associated with genetic diseases in human; B.R. is specialized in curation of vertebrate proteins; S.P. curates proteins across a variety of organisms.

The five UniProt curators first search PubTator for articles relevant to a specific protein (e.g. APC13 and Arabidopsis PubTator shows the exact same search results as PubMed except that the gene/protein names are highlighted in the titles. Once an article title is clicked in the search results, PubTator directs the users to its curation page (aka abstract page) where the automatic computer pre-annotations can be examined (and revised). Additional user comments can be further inserted in a text box when needed. All the edits and comments are recorded in PubTator and can be downloaded, either in bulk or by single article, for further analysis.

## Results

### Classification of the published literature

The selection of relevant and accurate literature is a key factor in the expert curation process (Poux, Magrane, et al., 2014). In UniProt, we do not aim to curate every available publication for a given protein. Instead, we concentrate curation efforts on publications that provide relevant novel information. For every paper included, there may be many other papers that are reviewed but excluded because they contain information that is redundant with existing literature-based annotations, provide weak evidence, are review articles or are simply beyond the scope of the information which UniProt captures. During the UniProtKB/Swiss-Prot literature triage process, evaluated articles can be classified into the following categories:

1. Curatable: For papers containing relevant information which are either selected for curation or are already present in UniProtKB/Swiss-Prot.
2. Not priority: For articles containing curatable information where the reported data are not considered high-priority so the papers were not selected for curation. For example, a paper reporting limited tissue specificity information in a low-priority organism, while microarrays databases provide more detailed and accurate information.
3. Not curatable: For articles that report information that is not relevant for inclusion in UniProtKB/Swiss-Prot. We further classify such articles into a number of subcategories:
  a. Not curatable - Out of scope: for papers that are not in the field of proteins or which use a protein as a marker.
  b. Not curatable – Redundant: for papers that describe relevant information which is already present in UniProtKB/Swiss-Prot as it has been previously reported in another reference.
  c. Not curatable - High-throughput: for articles that report high-throughput studies. As these studies generate a higher rate of false positives than classical assays, they are not selected for expert curation. Note that we have developed an expert-driven pipeline for inclusion of such publications that is distinct from expert curation (Breuza, Poux, et al., 2016).
  d. Not curatable – Insufficient evidence: for publications that report results that are not supported by strong experimental evidence and would need additional experimental confirmation for inclusion.
  e. Not curatable - Review/comment: for review and/or comment articles. Note that reviews that are not in the field of proteins were classified as ‘Not Curatable - Out of scope’.

Given these triage criteria, we performed a set of experiments using PubTator, a web-based text-mining tool in a PubMed-like environment, which provides access to all MEDLINE articles (Wei, Kao, et al., 2013a). A number of text-mining approaches have been integrated into PubTator to identify key biological entities such as gene, protein and species names, which are conveniently highlighted in the interface, facilitating the literature triage process. The interface has been adapted to suit the UniProt triage in a number of ways including linking of protein mentions to UniProt accessions, capability to filter by organism and ability to create collections of references. Most importantly, the tool enables the assignment of articles to the categories described above (Figure 1).

It is worth noting that the triage process involves a quick scan of the articles to identify the potential set for curation, and a more in-depth evaluation of that set to identify the curatable papers. The time spent for evaluating publications is variable: in many cases, the abstract is sufficient to determine if an article is relevant for curation and the evaluation is fast. In some cases, however, it is necessary to read the full-text article, which takes much more time. We used different approaches to evaluate the number of papers that are curatable.

### Proportion of PubMed which is curatable

We first addressed the question of the fraction of PubMed that is relevant for inclusion into UniProtKB/Swiss-Prot by examining a random selection of papers from PubMed. We triaged a random sample of 500 PubMed articles published from 2013 to 2015 (table 1). Our dataset is a representative set of PubMed as it has a similar overall distribution of articles based on indexed publication types in PubMed to the whole PubMed collection (Supplementary table 1). To ensure the accuracy of our classification method, the 500 articles were evaluated independently by 2 different curators and articles were assigned to the same category in more than 99% of the cases.

**Table 1.** Random sampling of PubMed

| Total | Curatable | Not<br>Priority | Not Curatable |  |  |  |  |
| --- | --- | --- | --- | --- | --- | --- | --- |
|  |  |  | Out of<br>scope | Redundant | High-throughput | Insufficient<br>evidence | Review/<br>comment |
| 500 | 19 | 17 | 452 | 2 | 2 | 5 | 3 |

Surprisingly, only 38 articles contained information potentially suitable for curation in UniProt (the ‘Curatable’, ‘Not priority’ and ‘Not Curatable – Redundant’ categories). Of these, only 19 articles contained relevant information for curation in UniProtKB/Swiss-Prot. However, when compared to existing information in the corresponding entries, only 10 (2%) would remain for full curation. An additional 9 papers provided interesting information but would not add essential knowledge or would provide weak evidence for an annotation. For example, Carry *et al.* describe a selective inhibitor of aurora kinases, while aurora kinase entries are well annotated and already describe selective inhibitors (UniProtKB 014965, Q96GD4 and 088445) (Carry, Clerc, et al., 2015). Similarly, Dong *et al.* describe expression profiling and results suggesting that TOR acts as a regulator of photosynthesis in *Arabidopsis thaliana* (UniProtKB Q9FR53) (Dong, Xiong, et al., 2015). Although the results are interesting, the experimental evidence is rather indirect and would require additional experimental support.

90% of the publications in this random set are completely outside the scope of UniProtKB.

### Number of curatable papers in a subset of journals

In a second approach, we assessed the number of articles that are curatable for UniProtKB/Swiss-Prot in a subset of journals (table 2). These journals (Cell, Developmental Cell, Elife, Genes and Development, Molecular Cell, Nature Cell Biology, Nature Genetics, Nature, PLoS Biology, PLoS Genetics, Science, The EMBO Journal, The Plant Cell) were selected based on their overall impact in the field combined with the fact that their content includes valuable information frequently prioritized for curation in UniProtKB/Swiss-Prot. PubTator automatically generates a weekly collection that includes the content of these journals. We have been using this tool to identify high-priority publications for curation for several years. During a six-month period, in addition to selecting articles for curation, we systematically classified all papers according to the criteria described in the previous section. We evaluated more than 5,000 publications from the subset of journals described above. Again, the proportion of articles that are curatable is quite low: only 13.1% of articles indexed in PubMed for these journals constitute high-priority targets for curation in UniProtKB/Swiss-Prot, while 65% are out of scope (table 2). Around 10% of articles evaluated concern either reviews or high-throughput studies.

**Table 2.** Monitoring articles in a collection of journals

| Total | Curatable | Not<br>Priority | Not Curatable |  |  |  |  |
| --- | --- | --- | --- | --- | --- | --- | --- |
|  |  |  | Out of<br>scope | Redundant | High-throughput | Insufficient<br>evidence | Review/<br>comment |
| 5013 | 659 | 681 | 3259 | 0 | 351 | 32 | 331 |

### Number of papers evaluated during the curation workflow

Finally, we tracked the number of publications that are evaluated during our routine protein-centric curation process, which typically starts with a literature search of a given protein. During a 6-month period, 5 curators systematically classified publications. To ensure that our analysis covered protein annotations from a variety of biological processes and a wide taxonomic range, we selected curators with different backgrounds and working in different annotation programs. During this period, these curators followed the classical UniProt expert curation workflow, a well-defined process that ensures that all records are handled in a consistent manner (read (Poux, Magrane, et al., 2014) for a more detailed description of the process), using PubTator to select and classify all publications evaluated during the curation process. In a 6-month period, more than 4,500 papers were evaluated by these 5 curators (table 3).

**Table 3.** Classification of papers evaluated during the expert curation process

| Total | Curatable | Not<br>Priority | Not Curatable |  |  |  |  |
| --- | --- | --- | --- | --- | --- | --- | --- |
|  |  |  | Out of<br>scope | Redundant | High-throughput | Insufficient<br>evidence | Review/<br>Comment |
| 4680 | 1398 | 641 | 1339 | 385 | 159 | 313 | 445 |

The proportion of papers that are curatable for UniProtKB/Swiss-Prot is low, even when specific terms such as gene or protein names are used to query the PubTator tool, as happens when curators perform searches during the curation process. Only 1,398 out of 4,680 articles evaluated (29.8%) were curatable, of which 584 were already present in UniProtKB/Swiss-Prot, meaning that only 814 new articles out of 4,680 (17%) were relevant for curation. It is important to note that, throughout the 6-month period of the study, both the proportion of curatable papers and the proportion of articles already present in UniProtKB/Swiss-Prot were stable, reinforcing the accuracy of our analysis (Supplemental table 2). Around a third of articles describe results that are out of the scope of UniProt, 10% of articles concern reviews and/or comment, and 8% of articles report redundant information, a frequent occurrence for papers reporting disease associations in human, where studies are made in different populations, generating a lot of redundant publications.

## Discussion

### A large fraction of papers indexed in PubMed are not relevant for UniProt curation

In UniProtKB/Swiss-Prot, knowledge is the driving factor for our expert curation effort and a targeted selection of papers is made in order to focus on publications that provide the maximum amount of high quality information. Curators assimilate all the information from various sources, reconcile any conflicting results and compile the data into a concise but comprehensive report, which provides a complete overview of the information available about a particular protein (for more information read (Poux, Magrane, et al., 2014)). Our literature triage clearly demonstrates that we evaluate a much higher number of articles than the 8,000-10,000 papers that we fully curate every year. Only 17% of articles evaluated during the curation process are curated in UniProtKB/Swiss-Prot. If we extrapolate these results to the entire expert curation team, we estimate that we read and evaluate between 50,000 and 70,000 articles every year. Measuring the number of curated publications is of course important, but it provides an incomplete picture of the complete set of papers evaluated during the curation process.

Our classification of articles during the triage process, however, should not be taken as a judgement on the quality of a publication. There can be different reasons why a publication is not selected for curation. Some excellent papers are not selected because the information relevant to UniProtKB is redundant with other publications present in the protein entry, while they contain outstanding information beyond the scope of UniProtKB. For example, Chen *et al.* developed an assay for measuring the deubiquitinase activity of OTULIN (Chen, Wang, et al., 2014) (all publications evaluated during this work are detailed in Supplemental table 6). The activity of OTULIN was already described in other publications whereas the development of the assay is not relevant for curation in UniProtKB. Moreover, our classification is not set in stone and can change: an article initially classified as ‘Not priority’ or ‘Not Curatable – Insufficient evidence’ may later be selected for curation when new data become available. A good example is provided by the DENND1B protein (UniProtKB Q6P3S1): an article reporting that variations in the DENND1B protein-coding gene are associated with susceptibility to asthma was not curated 4 years ago, because we considered that insufficient evidence was available at that time (Sleiman, Flory, et al., 2010). When the protein was updated in 2016, we revised our judgment based on new experimental results from other groups and reported this information from both articles (Sleiman, Flory, et al., 2010; Yang, Hojer, et al., 2016).

Our analysis also shows that a large fraction of the published literature is not curatable for UniProtKB/Swiss-Prot purposes. Scientific literature is highly redundant. While redundancy is extremely useful for reproducibility of results, a key challenge in science, we prioritize new knowledge over information already described in an entry. For example, Munch *et al.* reports a variant that is already described many times for human spastin entry (UniProtKB Q9UBP0) (Munch, Rolfs, et al., 2008). We read the paper but decided not to add it in UniProtKB/Swiss-Prot since it does not provide any additional information. In total, 77 articles reported redundant information for spastin. Many reviews and comments are also published: a total of 25 reviews were published for spastin. In many cases, we read such articles, but rarely integrate them because we favor curation of primary research results to allow for traceability of knowledge and so that curators can read the original research and make their own judgement on the data presented.

A large proportion of articles indexed in PubMed in the selection of journals that we parse every week are out of scope, even though these journals publish much valuable information for resources such as UniProtKB/Swiss-Prot. One reason for this is that articles that do not report experimental results, such as ‘news’ sections (Reardon, 2015) or corrections of previously published articles are all indexed in PubMed. Moreover, in general and multidisciplinary science journals, like Science and Nature, a lot of articles report on topics such as funding issues, political questions and climate change or publish articles not related to the life sciences (Hand, 2016). Last but not least, many articles describe new biological processes for which protein-coding genes have not yet been identified (Negishi, Miyazaki, et al., 2016; Zimmerman, Lin, et al., 2016).

When applied to all journals indexed in PubMed, the proportion of articles that are out of the scope of UniProtKB is more significant: the random sampling shows that 90% of articles indexed in PubMed in recent years are not relevant or suitable for curation in UniProtKB/Swiss-Prot. As an example, more than 15% of the publications found in PubMed are not written in English, meaning that they will not be curated even if they are within the scope of UniProtKB data. Moreover, many publications report biomedical studies such as response to medication or prevalence of a disease in different populations. Only a small proportion of journals indexed in PubMed are related to protein science. The low proportion of articles that are relevant for UniProt is confirmed by the number of journals cited in UniProt and PubMed: around 4,350 journals are cited in UniProt, a relatively small number compared to the 29,600 journals indexed in PubMed.

Even for papers that are in the field of proteins, a substantial proportion of articles is out of the scope of UniProtKB. For example, the protein RD29A in *Arabidopsis thaliana* (UniProtKB Q06738) is expressed in response to abiotic stress, such as salt, cold, abscisic acid and drought (Msanne, Lin, et al., 2011). Of 154 articles indexed for this protein, 70 were tagged as out of scope mostly because RD29A is used as a marker of abiotic stress. Another striking example is provided by the MKI67/Ki-67 protein (UniProtKB P46013). MKI67/Ki-67 is widely used as a marker of cell proliferation and constitutes the most widely used marker for comparing proliferation between tumor samples in the field of cancer research (Dowsett, Nielsen, et al., 2011; Richards-Taylor, Ewings, et al., 2016). More than 22,000 articles are indexed in PubMed concerning this protein, most of them using MKI67/Ki-67 as a marker. Ironically, while its function has been largely unclear for many years, recent results showed that its primary function is uncoupled from cell proliferation (Sobecki, Mrouj, et al., 2016). MKI67/Ki-67 is required to maintain dispersal of mitotic chromosomes by forming a steric and electrostatic charge barrier (Cuylen, Blaukopf, et al., 2016).

Note that we excluded proteins associated with thousands of articles, like MKI67/Ki-67, from the literature triage using PubTator, since proteins associated with thousands of articles of which only a small number are curatable would have provided strongly skewed results.

### Expert curation is sustainable

A major conclusion from the literature triage activity is that expert curation is sustainable. We estimate that we curate 35-45% of the proportion of PubMed that is relevant for UniProtKB/Swiss-Prot. The random sampling of PubMed showed that only 19 articles out of 500 contain curatable information, but that only 10 of them constitute high priority articles. Based on that, we estimate that a maximum of 2-3% of publications indexed in PubMed every year, between 20,000 and 25,000 articles, are curatable. This would suggest that we have accumulated a large backlog of curatable papers over the years, with a backlog of 12,000-15,000 papers for year 1, 24,000-30,000 papers for year 2, etc. This is however not the case: the curation workflow with a set of curators showed that 42% of the articles relevant for curation were already curated in UniProtKB/Swiss-Prot, while ‘only’ 58% concern new articles (Supplementary table 2). This demonstrates that we do not accumulate a large backlog. One reason is that knowledge captured from recently published articles replicates data from older papers and this makes up for knowledge that had been previously overlooked by rendering the older papers redundant for the purposes of UniProtKB curation.

A number of steps were taken in our study in order to reduce bias. A wide range of proteins were curated, coming from a number of different organisms, such as human, mouse, *Arabidopsis thaliana, Caenorhabditis elegans* and *Escherichia coli.* Moreover, proteins involved in a wide variety of biological processes were curated, including DNA repair pathways, circadian cycles, cytoskeleton regulation, chromatin regulation, embryonic development, and flowering. Finally, our analysis concerned proteins newly integrated into UniProtKB/Swiss-Prot as well as updates of records already present in UniProtKB/Swiss-Prot.

To ensure that we capture the maximum amount of curatable information and do not overlook important publications, we have put in place robust mechanisms to identify proteins to curate. Part of the prioritization is performed using PubTator, by parsing the tables of content of a number of journals, but other mechanisms are also used. For example, we track newly identified 3D-structures and changes in nomenclature in Model Organisms Databases (MODs) to identify newly characterized proteins. We also actively collaborate with other resources. For example, one of our curators is part of the Nomenclature Committee of the International Union of Biochemistry and Molecular Biology (IUBMB) and we actively participate in the creation of new EC numbers and their curation. Our users also frequently point out when information is incomplete. All these mechanisms ensure that we identify and curate relevant proteins and do not accumulate a backlog of curatable papers.

If we curate 35-45% of the relevant literature, what about the part that we do not capture? For vertebrate proteins, in particular human, we are more comprehensive and up-to-date: all proteincoding genes from human are present in UniProtKB/Swiss-Prot and the proportion of articles relevant for curation already curated in UniProtKB/Swiss-Prot is higher compared to other organisms (more than 50% for human). We of course overlook articles, but most important ones are captured or will be captured in the coming years. Curation efforts are focused on recently published articles with the largest numbers of papers added for the current year or the previous year. For a number of model organisms that are covered by the MODs, such as *Arabidopsis thaliana, Drosophila melanogaster* or *Caenorhabclitis elegans,* we are not yet complete, although we progress rapidly. We promote collaboration with other resources to exchange data and ensure that curation efforts are not duplicated, thus supplementing UniProt data with additional information curated by the MODs. It is also clear that the curation effort also reflects the size of research communities. We currently do not have sufficient resources to actively curate organisms studied by smaller scientific communities.

Our analysis however clearly shows that expert curation is sustainable and that the backlog of literature to curate is not as high as it first appears. It also suggests that a reasonable increase in funding would allow us to cover the vast majority of relevant publications. Similar conclusions were drawn recently by PomBase in an article that shows that the number of articles published on *Schizosaccharomyces pombe* has been stable over the years and that curation of this body of data is sustainable (Oliver, Lock, et al., 2016).

It is why the common belief that expert curation is highly expensive and time-consuming is incorrect. A recent article published by P. Karp demonstrated that the cost of curation is extremely modest compared to publication fees (Karp, 2016) and an independent survey assessing the value of biological database services concluded that the benefits to users and their funders are equivalent to more than 20 times the direct operational cost of the institute (http://www.ebi.ac.uk/about/news/press-releases/value-and-impact-of-the-european-bioinformatics-institute). From this perspective and from our analysis, expert curation should be considered as a major time-saver for the community for a very limited cost.

### The need for expert biocurators

If the increase in the number of papers published every year has no major impact on the scalability of expert curation, it however strongly affects the selection process, which is becoming a critical step in the curation process. A side effect of the increase in scientific publications concerns the growing presence of contradictory or incorrect results in the scientific literature. A number of articles have been published recently regarding the number of errors found in the scientific literature which are increasing to a level where science self-correction is no longer possible (Poux, Magrane, et al., 2014; Sarewitz, 2016). A recent article reported that more than 70% of researchers have tried and failed to reproduce another scientist’s experiments, and more than half have failed to reproduce their own experiments (Baker, 2016; Santori, 2016). The presence of erroneous or irreproducible results in the scientific literature highly complicates the task of users and can affect interpretation of data.

Besides erroneous data, we have to take into consideration that knowledge is a dynamic process and that our understanding of biology continues to evolve as new experiments confirm or contradict previous results. When new findings invalidate previous ones, old curation is revisited in the light of new knowledge and annotation from previous papers re-evaluated. This is where expert biocurators are indispensable in providing an overview of the latest data in the context of previous findings.

Experienced curators with a strong background in wet lab research are adept at dealing with conflicting or erroneous information. In UniProtKB/Swiss-Prot, a curator will read and curate a number of publications from different groups in different organisms, helping to resolve conflicting issues and providing a general overview of the state of research in the field. This helps to ensure maximal efficiency when curating groups of related proteins by providing the in-depth background knowledge required, thus reducing the time taken for curation of each individual protein. UniProtKB/Swiss-Prot entries also provide biological background and context: when information is in contradiction with previous reports, it is clearly mentioned in the entry (for more information, read (Poux, Magrane, et al., 2014)). For example, curation of the mitochondrial calcium uniporter in *Caenorhabditis elegans* (UniProtKB Q21121) following publication of its 3D-structure (Oxenoid, Dong, et al., 2016) generated a lot of collateral annotation: recent articles on the mitochondrial calcium uniporter were curated in different organisms, including human, mouse and *Dictyostelium discoideum* (UniProtKB Q8NE86, Q3UMR5, Q54LT0, respectively). Regulatory subunits of the mitochondrial calcium uniporter were also updated in order to provide a complete and up-to-date picture of the whole uniporter complex. This allowed the resolution of conflicting information, such as the topology of the SMDT1 regulatory subunit in human (UniProtKB Q9H4I9) and conflicting literature concerning MCUR1 (UniProtKB Q96AQ8). Full curation of articles, regardless of the species, is also essential in light of new technologies: researchers use different organisms for their research and jump from *Caenorhabditis elegans* to human or non-model organisms in the same paper. The article describing the 3D-structure of the mitochondrial calcium uniporter in *Caenorhabditis elegans* (Oxenoid, Dong, et al., 2016) also contained experiments performed with the mitochondrial calcium uniporter in human (UniProtKB Q8NE86).

The growth of scientific literature strongly suggests that expert curation is needed more than ever to separate the wheat from the chaff and select articles that provide the maximum amount of reliable information that users need.

### Improving efficiency of expert curation

Our results show that careful selection of papers is a critical step. The use of text-mining tools such as PubTator are of great help for curators by facilitating the literature triage process. Close collaboration between developers and curators is essential for the continued improvement of textmining accuracy and to save expert curation time. Moving forward, the data from the triage exercise performed in this work will be used to inform PubTator and tune the system to expedite the triage step in UniProt. Use of filters for excluding specific publication types (e.g. biographies), along with other methods, will be applied and evaluated. Collaboration represents a major challenge for integration of text-mining tools in the biocuration workflow and their customization and maintenance. Many of the tools are developed as part of a research proposal and/or for proof of concept, in isolation from their potential users and not with the end goal of adoption. There is an ongoing effort in BioCreative to promote interaction between the biocuration and text-mining communities to lower these barriers (Wang, S, et al., 2016). In fact, Pubtator is a product of the BioCreative 2012 challenge (Arighi, Carterette, et al., 2013).

In addition, other initiatives could reduce the burden of expert curation. Structuring knowledge in scientific publications would provide benefits, and initiatives such as SourceData (Liechti et al., submitted) or Forcell (Bandrowski, Brush, et al., 2015) are very promising. Marking up the content of articles with controlled vocabularies that can be read by a machine would facilitate extraction of data from literature sources and save curation time. This would also help in improving accuracy of normalization for text-mining and also help to map papers and feed the additional bibliography section of entries for users looking for publications that have not been curated yet or have not been selected for curation. The presence of a structured format will however not resolve all problems and will be a long process because it is likely that only a subset of journals will adopt a structured format initially with the remainder containing free text information for an extended period. More importantly, the major challenge for users will still be the identification and selection of appropriate data and the extraction of reliable information.

The problem would be the same if authors curated their own publications. There would still be a need for expert curators to extract the real knowledge from the ocean of data. The example of the MKI67/Ki-67 protein described before is striking, with thousands of papers claiming that this protein is involved in cell proliferation compared to only a subset of articles that show that this is not the case. This case may seem extreme, but there are many examples like this and we encountered much contradictory or conflicting information in the publications evaluated and curated during this analysis, especially in the biomedical fields. Mechanisms will have to be found to filter knowledge and biocurators will continue to play a key role in this process. For these different reasons, we strongly believe that expertly curated databases will remain the cornerstone of scientific research, allowing users to keep up with the generation and evolution of knowledge and to provide goldstandard information.

## Acknowledgement

UniProt has been prepared by Maria Jesus Martin, Claire O’Donovan, Emanuele Alpi, Ricardo Antunes, Benoit Bely, Mark Bingley, Carlos Bonilla, Ramona Britto, Borisas Bursteinas, Andrew Cowley, Alan Da Silva, Maurizio De Giorgi, Tunca Dogan, Francesco Fazzini, Leyla Garcia Castro, Luis Figueira, Penelope Garmiri, George Georghiou, Daniel Gonzalez, Emma Hatton-Ellis, Weizhong Li, Wudong Liu, Rodrigo Lopez, Jie Luo, Yvonne Lussi, Alistair MacDougall, Andrew Nightingale, Barbara Palka, Klemens Pichler, Diego Poggioli, Sangya Pundir, Luis Pureza, Guoying Qi, Steven Rosanoff, Rabie Saidi, Tony Sawford, Aleksandra Shypitsyna, Elena Speretta, Edward Turner, Nidhi Tyagi, Vladimir Volynkin, Tony Wardell, Kate Warner, Xavier Watkins, Rossana Zaru and Hermann Zellner at the European Bioinformatics Institute; Ioannis Xenarios, Lydie Bougueleret, Alan Bridge, Nicole Redaschi, Lucila Aimo, Ghislaine Argoud-Puy, Andrea Auchincloss, Kristian Axelsen, Parit Bansal, Delphine Baratin, Marie-Claude Blatter, Brigitte Boeckmann, Jerven Bolleman, Lionel Breuza, Cristina Casal-Casas, Edouard de Castro, Elisabeth Coudert, Beatrice Cuche, Mikael Doche, Dolnide Dornevil, Severine Duvaud, Anne Estreicher, Marc Feuermann, Elisabeth Gasteiger, Sebastien Gehant, Vivienne Gerritsen, Arnaud Gos, Nadine Gruaz-Gumowski, Ursula Hinz, Chantal Hulo, Florence Jungo, Guillaume Keller, Vicente Lara, Philippe Lemercier, Damien Lieberherr, Thierry Lombardot, Xavier Martin, Patrick Masson, Anne Morgat, Teresa Neto, Nevila Nouspikel, Salvo Paesano, Ivo Pedruzzi, Sandrine Pilbout, Monica Pozzato, Manuela Pruess, Catherine Rivoire, Michel Schneider, Christian Sigrist, Karin Sonesson, Sylvie Staehli, Andre Stutz, Shyamala Sundaram, Michael Tognolli, Laure Verbregue and Anne-Lise Veuthey at the SIB Swiss Institute of Bioinformatics; Cathy H. Wu, Leslie Arminski, Chuming Chen, Yongxing Chen, John S. Garavelli, Hongzhan Huang, Kati Laiho, Peter McGarvey, Darren A. Natale, Karen Ross, C. R. Vinayaka, Qinghua Wang, Yuqi Wang, Lai-Su Yeh and Jian Zhang at the Protein Information Resource.

